# Single-dose VSV-Sudan virus vaccine protects from lethal Sudan virus infection within one week: a challenge study in macaques

**DOI:** 10.1101/2025.03.26.645555

**Authors:** Paige Fletcher, Kyle L. O’Donnell, Friederike Feldmann, Joseph F. Rhoderick, Chad S. Clancy, Cecilia A. Prator, Brian J. Smith, Bronwyn M. Gunn, Heinz Feldmann, Andrea Marzi

**Affiliations:** Laboratory of Virology, Division of Intramural Research, National Institute of Allergy and Infectious Diseases, National Institutes of Health, Rocky Mountain Laboratories, Hamilton, MT 59840, USA; Rocky Mountain Veterinary Branch, Division of Intramural Research, National Institute of Allergy and Infectious Diseases, National Institutes of Health, Rocky Mountain Laboratories, Hamilton, MT 59840, USA; Paul G. Allen School for Global Health, Washington State University, Pullman, WA, USA

**Keywords:** Filovirus, VSV-SUDV, VSV-EBOV, nonhuman primate, cross-protection

## Abstract

**Background:** The Sudan virus (SUDV) outbreaks in recent years including the ongoing outbreak in Uganda created a public health emergency beyond the Eastern Africa region. Currently, there are licensed countermeasures for Ebola virus (EBOV); however, there are no licensed vaccines or therapeutics against SUDV.

**Methods:** We developed a vesicular stomatitis virus (VSV)-based vaccine expressing the SUDV glycoprotein. Cynomolgus macaques were vaccinated intramuscularly with a single dose of VSV-SUDV either one month or one week prior to SUDV challenge. A third group was vaccinated with a single dose of VSV-EBOV one month prior to SUDV challenge to assess its cross-protective potential, and a control group received an unrelated VSV-based vaccine.

**Results:** All vaccinated nonhuman primates (NHPs) developed antigen-specific IgG within 2 weeks of vaccination, including cross-reactive responses. After challenge with a lethal dose of SUDV, all VSV-SUDV-vaccinated NHPs were uniformly protected from disease. In contrast, the VSV-EBOV-vaccinated and control NHPs succumbed to disease between day 5 and 7 after challenge presenting with classical signs of Sudan virus disease associated with high titer viremia, high viral organ load, dysregulated cytokine profiles and typical pathological changes. The humoral immune response in the NHPs vaccinated with VSV-SUDV one month before challenge resulted in a profound and sustained antibody response with a diverse functionality profile which was not observed to the same extend in NHPs vaccinated one week before challenge.

**Interpretation:** We demonstrated that a single dose of VSV-SUDV protected NHPs from lethal SUDV infection within one week. The fast-acting nature makes VSV-SUDV an ideal countermeasure for ring vaccination during outbreaks of Sudan virus disease. In contrast, VSV-EBOV provided no relevant protection against SUDV infection in NHPs highlighting the need for species-specific filovirus vaccines.

## Introduction

Filoviruses continue to pose a significant threat to regional public health and beyond, particularly in their endemic areas in Africa ^1^. In the last decade alone, Ebola virus (EBOV) caused the West African epidemic and several large outbreaks in the Democratic Republic of Congo resulting in over 30,000 cases and 12,000 deaths ^2^. In addition, Marburg virus (MARV), Sudan virus (SUDV) and Bundibugyo virus have caused outbreaks albeit smaller in West, Central and East Africa highlighting the continued circulation of filoviruses in their respective reservoirs with expanding endemic areas ^2,3^.

SUDV is the second most important filovirus for public health in Africa; 9 outbreaks have occurred mainly in Sudan and Uganda with the largest on record in Gulu, Uganda causing 425 infections with 224 fatalities ^2^. More recently, an outbreak of Sudan virus disease (SVD) in Uganda lasted from September 2022 until January 2023 and resulted in a total of 164 cases and 77 fatalities ^4^. Currently, Uganda is again experiencing an ongoing SUDV outbreak in its capital, Kampala ^5^. There are no approved vaccine or treatment options available for SUDV; however, a vesicular stomatitis virus (VSV)-based vaccine expressing the SUDV glycoprotein (GP), termed VSV-SUDV, is currently being used in a clinical trial in Uganda ^6^. In addition, a monoclonal antibody specific for SUDV GP as well as the antiviral remdesivir are being administered to infected people ^5^.

SUDV forms a separate species in the *Ebolavirus* genus, family *Filoviridae* ^7^. Previous studies demonstrated that surviving nonhuman primates (NHPs) from EBOV infections generally succumb to a subsequent SUDV challenge and *vice versa* indicating limited cross-protection between those ebolavirus species ^8,9^. Thus, licensed EBOV vaccines and direct-acting specific therapeutics like monoclonal antibodies are unlikely to provide protection against SUDV infection. Previously we demonstrated that VSV-SUDV is uniformly protective within 28 days when a single high dose was administered to NHPs with preexisting immunity to VSV-EBOV ^9^. These results add to the growing body of evidence that pre-existing vector immunity has only limited impact on VSV-filovirus vaccine efficacy ^9–11^. However, only limited data is available on the cross-protective potential of species-specific VSV-filovirus vaccines in NHP models ^12–14^.

Therefore, we determined the protective efficacy of a single high dose VSV-SUDV or VSV-EBOV vaccination against lethal SUDV challenge. In addition, we investigated if the VSV-SUDV vaccine possesses fast-acting potential to be used to control Sudan virus disease (SVD) outbreaks ^15,16^.

## Results

### VSV-SUDV vaccination protects NHPs from SVD

NHPs (n=6 per group) were vaccinated intramuscularly (IM) 28 days prior to challenge (d-28) with 1x 10^7^ plaque-forming units (PFU) of either VSV-SUDV, VSV-EBOV or a control vaccine (VSV-LASV). Another group was vaccinated with 1x 10^7^ PFU VSV-SUDV on d-7. Vaccine viremia was short-lived in all vaccinated NHPs regardless of the vaccine construct peaking on d1 and resolving within a week after vaccination (Fig. 1A). On d0, all 24 NHPs were challenged IM with a lethal dose of 1×10^4^ median tissue culture infectious doses (TCID_50_) of SUDV. Clinical signs of disease were assessed at least once daily during the acute disease phase by assigning a score for general appearance, skin and fur, nose/mouth/eyes/head, respiration, feces and urine, food intake, and locomotor activity (Fig. 1B). NHPs in the VSV-EBOV and the control groups started to display signs of disease on d3. Unexpectedly, disease in the VSV-EBOV group progressed significantly faster compared to the control group leading to euthanasia of the NHPs on d5-6 (Fig. 1C). NHPs in the control group reached endpoint criteria on d6-7 and were euthanized. All 12 NHPs euthanized between d5-7 presented with high levels of SUDV RNA in the blood (Fig. 1D) which corresponded with high concentrations of soluble SUDV glycoprotein (sGP) in the serum (Fig. 1E). All non-protected animals displayed a decrease in thrombocytes (Fig. 1F), decreases in white blood cell (WBC), specifically neutrophil and lymphocyte counts (fig. S1A-C), elevated liver (Fig. 1G; Fig S1D,E) and kidney (Fig. 1H; Fig. S1F) enzyme levels as well as a decrease in serum albumin, total protein, and calcium (Fig. S1G-I), all characteristic parameters for SVD in NHPs. The VSV-EBOV group demonstrated with a stress leukogram as evidenced by increases in WBCs and neutrophils on d3 together with a decrease of lymphocytes. The control NHPs did not develop this immune cell signature. In contrast, NHPs vaccinated with VSV-SUDV at either d-28 or d-7 survived the lethal challenge with minimal signs of disease and presented with very low and temporary SUDV RNA levels in the blood on d6 and d9 (Fig. 1D).

**Figure 1.**
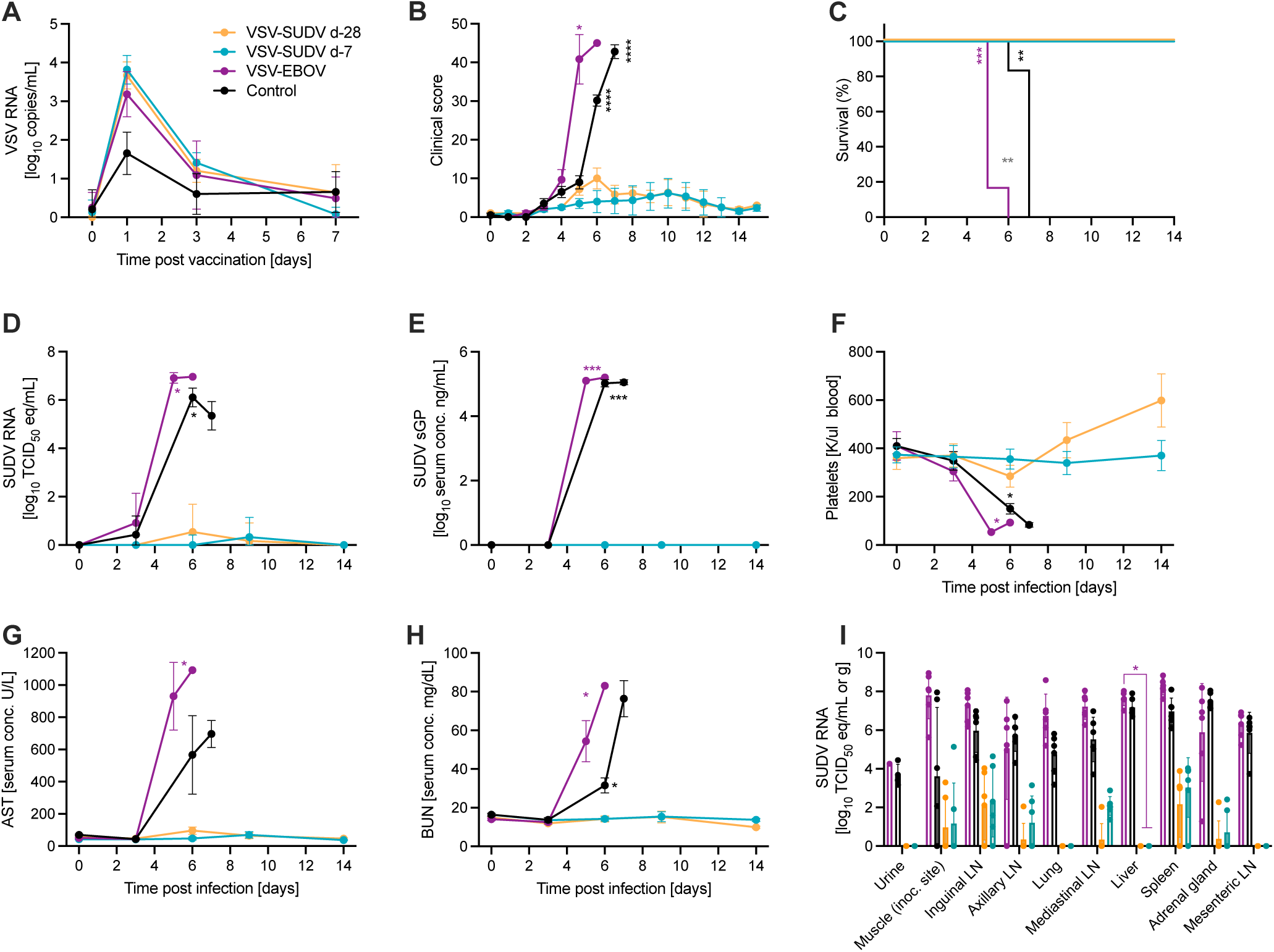
Survival and clinical changes in NHPs after SUDV challenge. NHPs (n=6 per group) were vaccinated 28 days before challenge (d-28) with either VSV-SUDV, VSV-EBOV or control vaccine (VSV-LASV). Another group was vaccinated with VSV-SUDV 7 days before challenge (d-7). On day 0, all 24 NHPs were challenged with a lethal dose of SUDV. (A) VSV RNA in the blood after vaccination. (B) Clinical scores, (C) survival, (D) viremia by RT-qPCR, and (E) SUDV sGP levels in serum after challenge are shown. (F) Platelet count in whole blood samples after challenge. Serum levels of (G) aspartate aminotransferase (AST), and (H) blood urea nitrogen (BUN) were determined. (I) SUDV RNA levels in tissue samples collected at necropsy. Geometric mean and geometric SD are depicted in A, D, E, I. Mean and standard error of the mean are shown in B, C, F-H. Significant differences in the survival curves were determined performing Log-Rank analysis. All other data were evaluated for statistical significance using two-way ANOVA with Tukey’s multiple comparisons. Statistical significance is indicated as \**p*<0.05, \*\**p*<0.01, \*\*\**p*<0.001, and \*\*\*\**p*<0.0001.

### Non-protected NHPs developed SVD with interstitial pneumonia

For all non-protected animals, necropsies were performed when euthanasia criteria were met between d5-7 and samples for virologic and histopathologic analysis were collected. SUDV RNA was detected at high levels in tissue samples collected at necropsy from these NHPs (Fig. 1I). Histologic lesions consistent with classic SUDV disease were observed in the liver and spleen in both the control and VSV-EBOV-vaccinated NHPs (Fig. 2). Mild to severe lymphoid necrosis with lymphoid depletion was observed in the spleen of every NHP (Fig. 2A, D). Immunohistochemical evaluation showed follicular presence of SUDV antigen in sections of spleen of all control and VSV-EBOV-vaccinated NHPs (Fig. 2G, J). Hepatic lesions included single cell hepatocellular necrosis, random or centrilobular coagulative necrosis and sinusoidal fibrin thrombi disseminated throughout evaluated sections (Fig. 2B, E). Minimal to mild lymphohistiocytic periportal hepatitis was observed in all control NHPs (Fig. 2E). Additionally, Kupffer cells and some hepatocytes showed SUDV antigen in these groups (Fig. 2H, K). A prominent lesion observed in at least one lung lobe of every control or VSV-EBOV-vaccinated NHPs was pyogranulomatous vasculitis extending into interstitial pneumonia (Fig. 2C, F).

**Figure 2.**
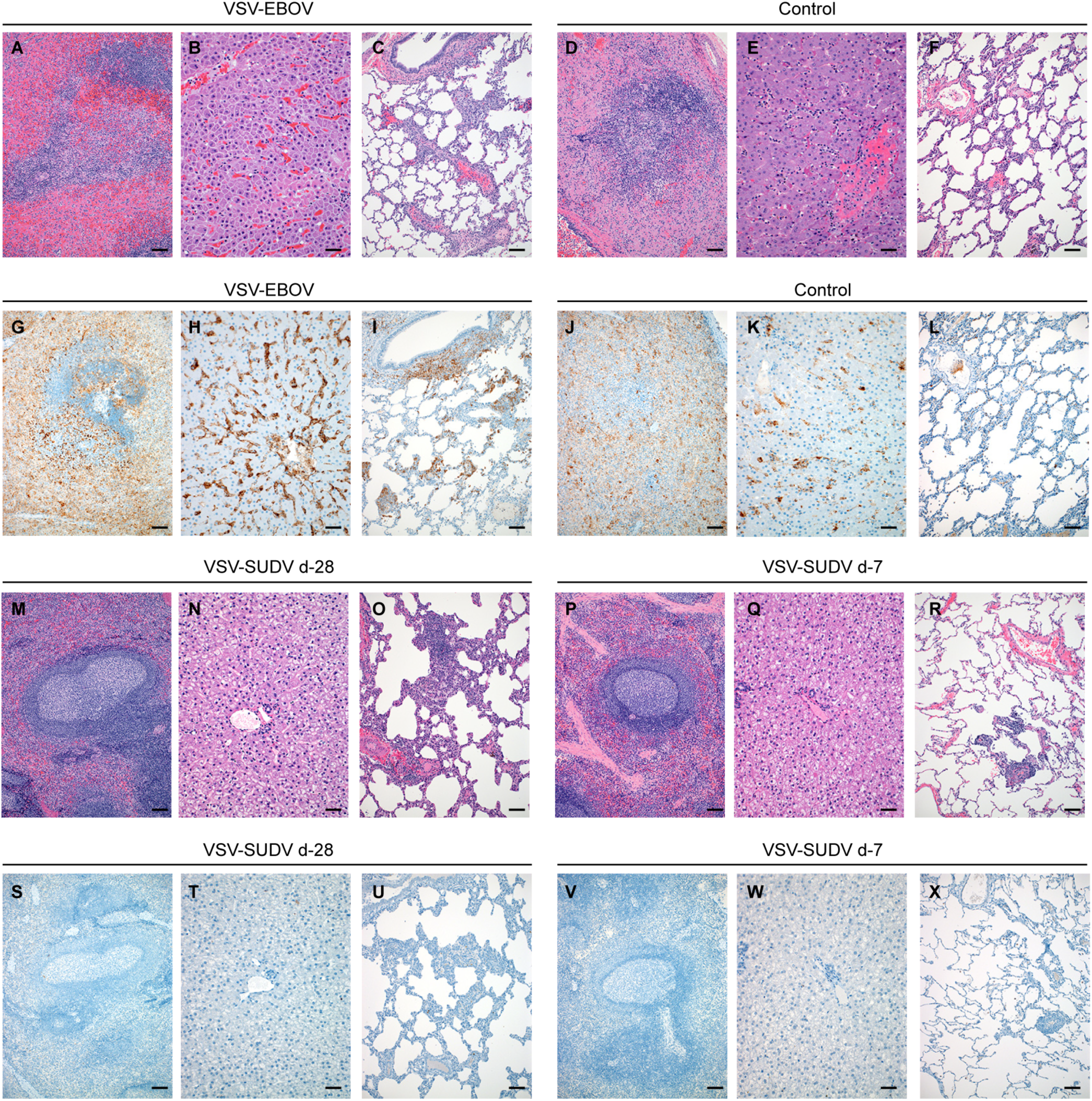
Histopathologic and immunohistochemical findings in NHPs after SUDV challenge. (A) Spleen exhibiting follicular lymphoid depletion and neutrophilic splenitis (hematoxylin and eosin (H&E), 100x). (B) Liver exhibiting random coagulative necrosis, sinusoidal fibrin thrombi and individual hepatocellular necrosis (H&E, 200x). (C) Lung with moderate pyogranulomatous vasculitis and interstitial pneumonia (H&E, 100x). (D) Spleen exhibiting lymphoid depletion and follicular lymphoid necrosis (H&E, 100x). (E) Liver exhibiting hepatocellular necrosis and neutrophilic hepatitis (H&E, 200x). (F) Lung exhibiting mild vasculitis extending into interstitial pneumonia (H&E, 100x). (G) Spleen section with immunoreactivity noted in both red and white pulp (immunohistochemistry (IHC), 100x). (H) Liver section showing nearly ubiquitous Kupfer cell and endothelial cell with rare hepatocyte immunoreactivity (IHC, 100x). (I) Lung section with strong immunoreactivity visible in pericytes, smooth muscle cells, foci of interstitial inflammation and pulmonary macrophages (IHC, 100x). (J) Spleen section with immunoreactivity noted in both red and white pulp (IHC, 100x). (K) Liver section showing immunoreactivity largely within endothelial cells and Kupfer cells (IHC, 200x). (L) Rare immunoreactivity is limited to foci of interstitial pneumonia and pulmonary macrophages (IHC, 100x). (M) Spleen with a large, reactive lymphoid follicle characterized by a prominent germinal center and distinct and well-defined mantle zone (H&E, 100x). (N) Liver exhibiting no significant histologic changes (H&E, 200x). (O) Lung showing a focus of moderate, chronic interstitial pneumonia with a prominent aggregate of lymphocytes and infiltration of foamy macrophages within alveolar spaces (H&E, 100x). (P) Spleen with a large, reactive lymphoid follicle characterized by a prominent germinal center and distinct and well-defined mantle zone (H&E, 100x). (Q) Liver exhibiting no significant histologic changes (H&E, 200x). (R) Lung exhibiting a small aggregate of perivascular lymphocytes that spill into adjacent alveolar spaces (H&E, 100x). (S) Spleen (100x) and (T) liver (200x) sections showing lack of immunoreactivity. (U) Lung section showing lack of immunoreactivity within a focus of interstitial pneumonia (IHC, 100x). (V) Spleen (100x) and (W) liver (200x) sections showing lack of immunoreactivity. (X) Lung section showing lack of immunoreactivity within a focus of interstitial pneumonia (IHC, 100x). Scale bar represents 100 μm.

Interesting, pyogranulomatous inflammation was categorized as moderate in at least one lung lobe of n=5/6 of VSV-EBOV-vaccinated NHPs whereas moderate inflammation was only detected in the lung lobes of n=1/6 control NHPs. Prominent fibrin thrombi and thrombo-emboli ranging from capillary to large caliber arteries were observed in at least one evaluated lung lobe in each of the control or VSV-EBOV-vaccinated NHPs. Evaluation of lung sections showed SUDV antigen in pneumocytes, pulmonary macrophages, endothelial cells, and smooth muscle cells of at least one evaluated section of lung in all control and VSV-EBOV-vaccinated NHPs (Fig. 2I, L). Additional classic histologic changes observed in control and VSV-EBOV-vaccinated NHPs include lymphohistiocytic duodenitis associated with hemorrhage and edema.

In VSV-SUDV-vaccinated NHPs at study end (d42), the only histologic changes observed in peripheral nodes and sections of the spleen included lymphoid hyperplasia and sinus histiocytosis (Fig. 2M, P) consistent with a response to remaining low levels of SUDV RNA (Fig. 1I). Evaluation of lung sections of VSV-SUDV-vaccinated NHPs revealed lymphohistiocytic vasculitis extending into alveolar septa (Fig. 2O, R). No significant histopathologic lesions were observed in the duodenum of any VSV-SUDV-vaccinated NHP. SUDV antigen was not observed in evaluated sections of lymph node, spleen, liver, or lung in any VSV-SUDV-vaccinated NHPs (Fig. 2S-X).

### Only non-protected NHPs developed a cytokine storm

Filovirus disease, including SVD, is associated with a dysregulated cytokine response ^17^, therefore, serum cytokine and chemokine responses in NHPs after vaccination and challenge were assessed. Serum concentrations of GM-CSF, MCP-1, IL-10, IL-12p70, IFN-α2a, IFNγ, IL-4, IL-6, IL-15, IP-10, IL-1β, and TNFα were measured for one week after vaccination and for 2 weeks after challenge. After vaccination all groups had a similar response to the replicating viral vaccine (Fig. S2). Only the control NHPs developed a significant increase of MCP-1 one day after vaccination. After challenge, only the VSV-EBOV and control NHPs developed elevated cytokine and chemokine levels indicative of a cytokine storm (Fig. 3). Levels of MCP-1, IL-15, and IP-10 were significantly increased in the serum of NHPs during end-stage SVD. In contrast, all VSV-SUDV-vaccinated NHPs maintained normal cytokine levels up to d14 after challenge (Fig. 3).

**Figure 3.**
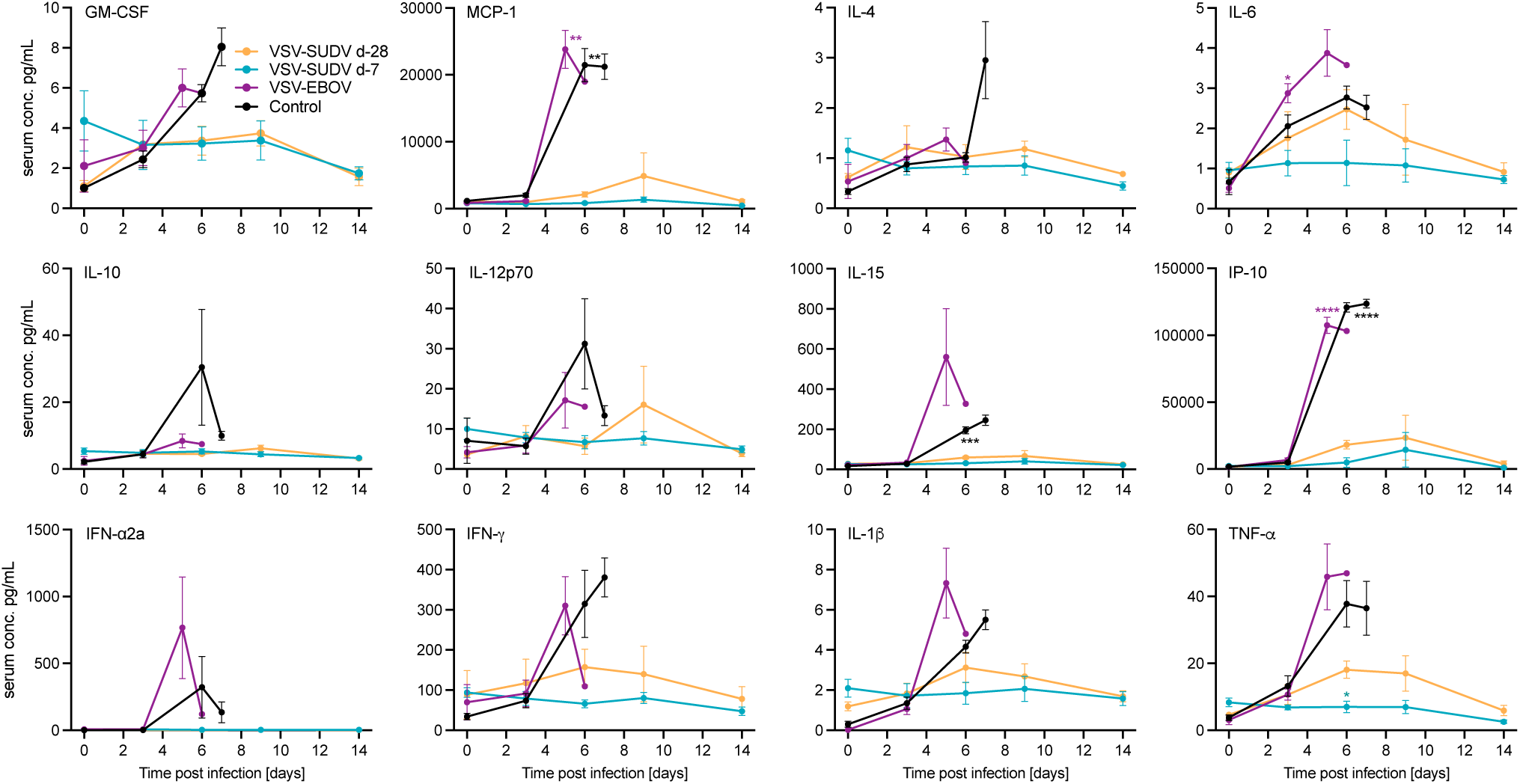
Cytokine and chemokine responses after challenge. Expression levels of selected cytokines or chemokines were determined after SUDV-challenge. Mean and standard error of the mean are depicted. Statistical significance was determined by two-way ANOVA with Tukey’s multiple comparisons. Statistical significance is indicated as \*\**p*<0.01, \*\*\**p*<0.001, and \*\*\*\**p*<0.0001.

Since significant lung vaso-centric changes in NHPs succumbing to SVD were observed during necropsy and on histopathology, cytokine and chemokine expression levels as well as the amount of SUDV sGP present in lung samples of all NHPs collected at the time of euthanasia were assessed. Only lung samples from NHPs in the VSV-EBOV and control groups had significantly higher cytokine and chemokine expression profiles. Notably, GM-CSF, TNFα, IL-6, IP-10, and MCP-1 were only significantly upregulated in lung samples from the VSV-EBOV group compared to both VSV-SUDV groups but not compared to the control NHPs (Fig. 4A).

**Figure 4.**
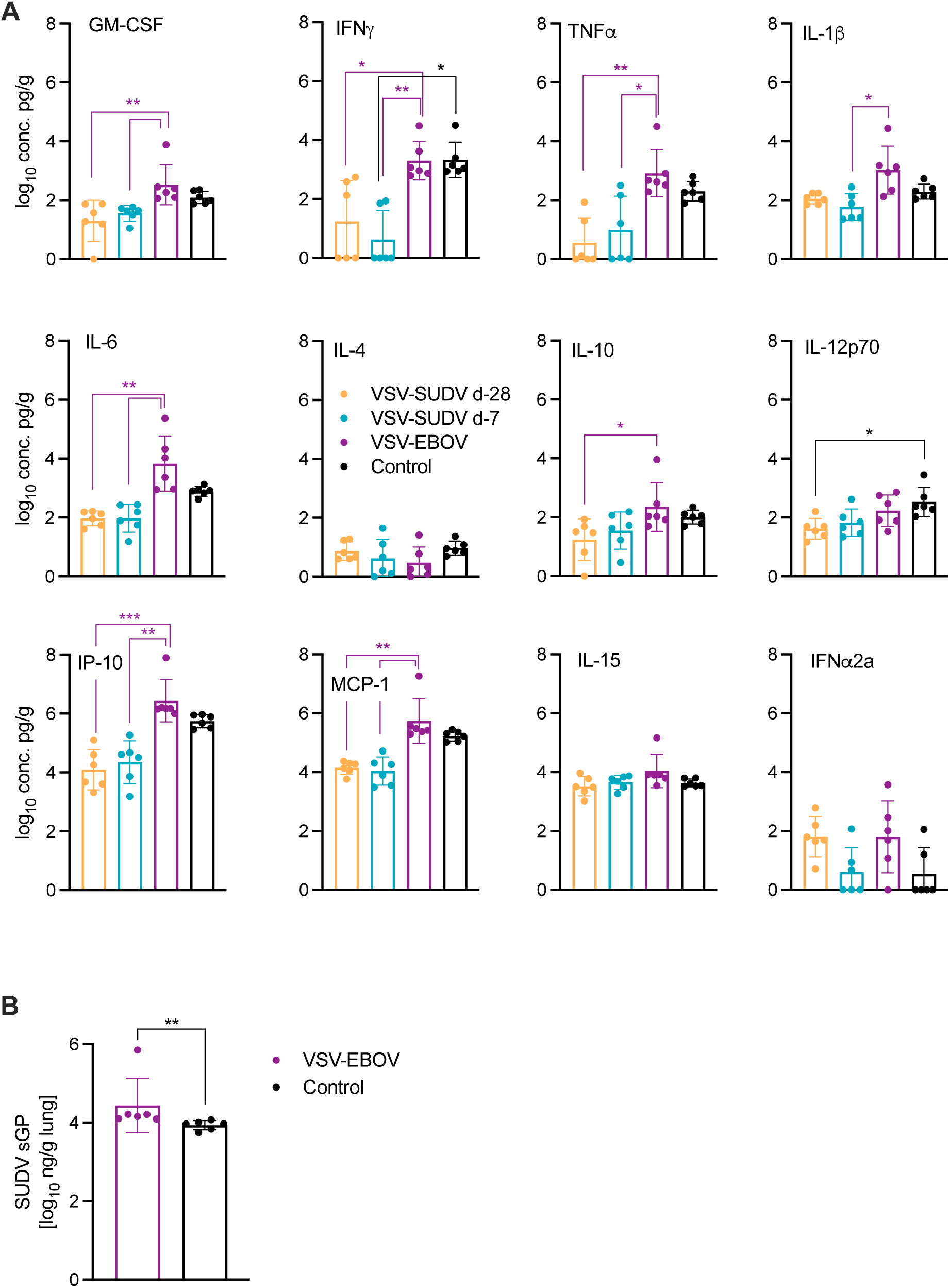
Levels of cytokines, chemokines and SUDV sGP in the lungs of NHPs. (A) Expression levels of selected cytokines or chemokines were determined in lung samples from NHPs collected at the time of necropsy. Geometric mean and geometric SD are depicted. Statistical significance was determined by Kruskal-Wallis test with Dunn’s multiple comparisons. (B) SUDV sGP levels in NHP lungs. Geometric mean and geometric SD are depicted. Statistical significance was determined by Mann-Whitney test. Statistical significance is indicated as \**p*<0.05, \*\**p*<0.01, and \*\*\**p*<0.001.

While there wasn’t a significant difference in the lung cytokine expression profile or SUDV lung RNA levels (Fig. 1I) between the VSV-EBOV and control groups, SUDV sGP levels were significantly higher in the VSV-EBOV group compared to control NHPs (Fig. 4B). This difference was not observed in serum SUDV sGP levels (Fig. 1E). All VSV-SUDV-vaccinated NHPs presented with normal cytokine levels in the lungs at the study end (d42) and since there was no indication of SUDV viremia or SUDV sGP in the serum during the acute disease phase (Fig. 1D, E), SUDV sGP levels were not assessed as they correlate with viral RNA levels ^18^.

### Protected NHPs developed robust antigen-specific immune responses

The total antigen-specific IgG response has been identified as the main mediator of protection for VSV-filovirus vaccines ^19,20^. Therefore, SUDV GP-specific IgG serum levels were determined longitudinally from d-28 to d42 and as expected, revealed robust responses for the VSV-SUDV d-28 group at the time of challenge (Fig. 5A). The boosting effect of the challenge virus was not as substantial for the VSV-SUDV d-7 group, presumably because the shorter time between vaccination and challenge. However, for both groups the exposure to the challenge virus boosted the SUDV GP-specific response (Fig. 5A). The VSV-EBOV-vaccinated NHPs developed intermediate levels of IgG cross-reactive with SUDV GP which decreased after challenge likely due to IgG consumption from SUDV replication (Fig. 5A). In contrast, these NHPs developed significant titers of EBOV GP-specific IgG (Fig. S3A). The d-28 VSV-SUDV-vaccinated NHPs also developed cross-reactive IgG albeit at lower levels (Fig. S3A). Analysis of serum samples from the control NHPs did not result in significant levels of EBOV GP-or SUDV GP-specific IgG (Fig. 5A; Fig. S3A). A recent publication highlighted the importance of EBOV sGP-specific antibodies for durable protection in NHPs ^21^. When we compared the IgG titers specific to SUDV sGP in serum at selected time points between the groups we found that the levels were associated with the serum levels of SUDV GP-specific IgG (Fig. 5A, B). This was not the case for EBOV sGP-specific IgG where levels remained constant throughout the study in surviving NHPs but EBOV GP-specific IgG levels declined over the duration of the study (Fig. S3B).

**Figure 5.**
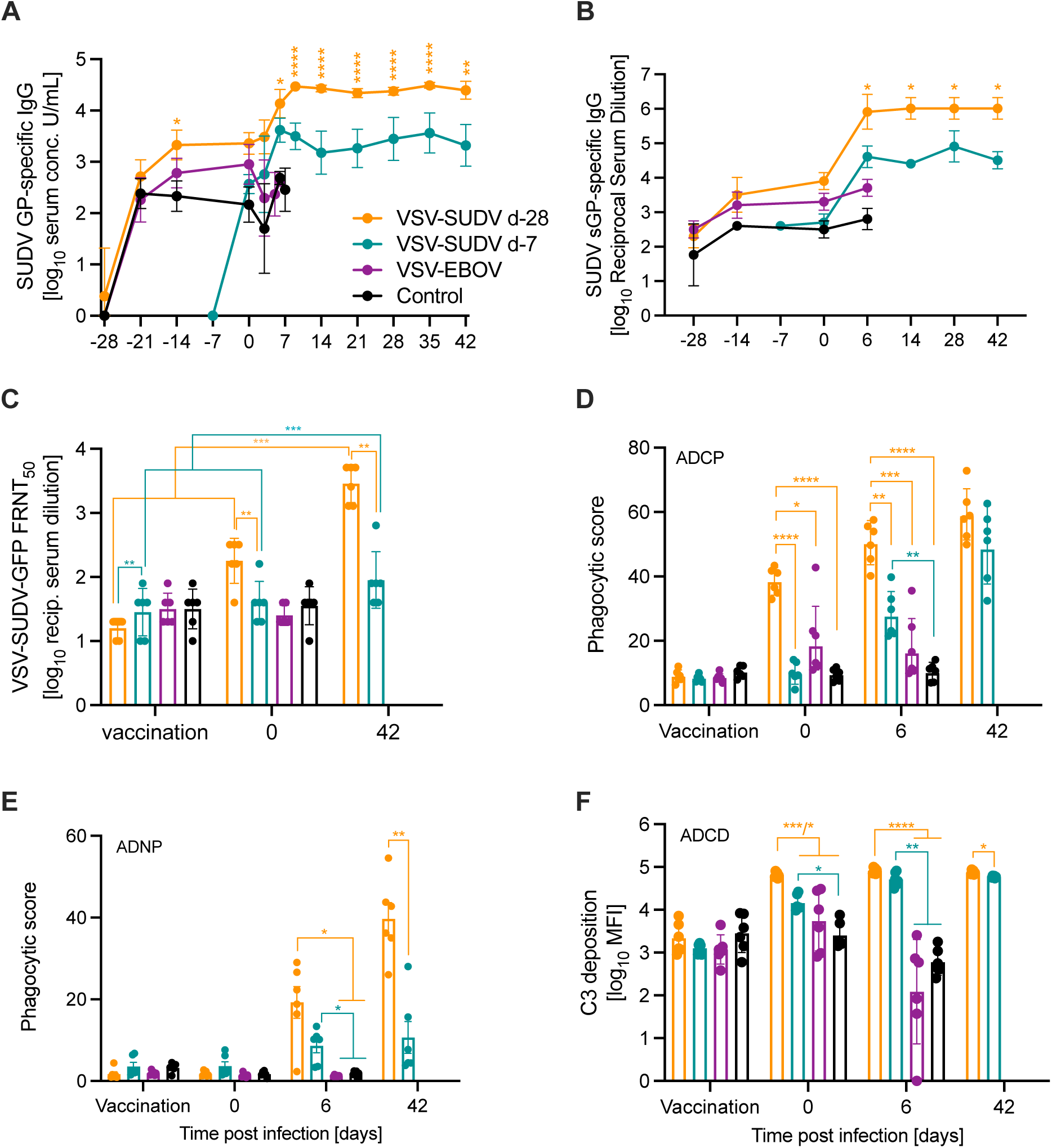
Humoral immune responses after vaccination and SUDV challenge. (A) SUDV GP-specific and (B) SUDV sGP-specific serum IgG levels were determined over time. (C) Serum neutralization presented as 50% fluorescence reduction (FRNT_50_) of GFP-positive cells at the time of vaccination (day - 28 or day −7, respectively), challenge (day 0) and study end (day 42). (D) Antibody-dependent cellular phagocytosis (ADCP), (E) antibody-dependent neutrophil phagocytosis (ADNP), and (F) antibody-dependent complement deposition (ADCD) were determined from serum. Geometric mean and geometric SD are depicted in A-C, F. Mean and standard error of the mean are shown in D, E. Statistical significance was determined by two-way ANOVA with Tukey’s multiple comparisons (4 groups) or Sidak’s multiple comparisons (2 groups). Statistical significance is indicated as \**p*<0.05, \*\**p*<0.01, \*\*\**p*<0.001, and \*\*\*\**p*<0.0001.

Next, serum neutralization titers were evaluated using a previously established VSV-SUDV-GFP neutralization assay ^9^ with results presented as 50% fluorescence reduction (FRNT_50_) of GFP-positive cells at the time of vaccination (d-28 or d −7), challenge (d0) and study end (d42). Only VSV-SUDV d-28-vacinated NHPs had developed a significant neutralizing response at the time of challenge which was boosted by the SUDV challenge (Fig. 5C). In contrast, the neutralizing antibody titers in the d-7 group remained limited throughout the study but showed a significant increase on d42 (Fig. 5C). NHPs in the control as well as VSV-EBOV-vaccinated group developed no neutralizing response (Fig. 5C).

Because the VSV-EBOV group developed accelerated disease and succumbed significantly earlier than the control group, we investigated if the EBOV GP-specific immunity which is unspecific to the challenge virus could have resulted in antibody-dependent enhancement (ADE) of SUDV infection as there is *in vitro* evidence for ADE in EBOV infection ^22–24^. We compared serum samples from the time of vaccination (d-28) and the time of challenge (d0) of the VSV-EBOV and control groups in this assay using SUDV-GFP. Serum samples in both groups did not reach levels associated with ADE described by others for filoviruses (Fig. S3C) ^24^.

In addition to the antibody response analysis described above alternative antibody Fc-effector functions were investigated, specifically the antibody-dependent cellular phagocytosis (ADCP), antibody-dependent complement deposition (ADCD), antibody-dependent neutrophil phagocytosis (ADNP), and antibody-dependent natural killer cell activation (ADNKA). NHPs vaccinated with VSV-SUDV at d-28 demonstrated significantly more ADCP at the time of challenge which continued through d6 (Fig. 5D). Taking a closer look at the cell type-specific phagocytosis we investigated whether neutrophils would facilitate phagocytosis. NHPs vaccinated at either time point with VSV-SUDV demonstrated significantly higher ADNP levels on d6 when the control and VSV-EBOV cohorts succumbed to disease (Fig. 5E). At study end (d42), the levels remained steady in the VSV-SUDV d-7 group but had further increased in the d-28 VSV-SUDV group (Fig. 5E). In the same manner, NHPs vaccinated with VSV-SUDV at either time point showed a significantly stronger induction of ADCD at the time of challenge and on d6 (Fig. 5F). Interestingly, at the time of challenge, ADCD was significantly increased in the VSV-SUDV d-28 group compared to the cohort vaccinated on d-7 with VSV-SUDV (Fig. 5F). The final functionality assessed was ADNKA (Fig. S3D-F) which yielded little differences with only the control group demonstrating a significantly lower amount of IFNγ induction at the time of challenge (Fig. S3E).

### Limited T cell responses in vaccinated and challenged NHPs

Despite the limited impact of T cell responses on protective efficacy of VSV-filovirus vaccines ^19,20,25^, CD4 T cell responses after vaccination and challenge were analyzed to evaluate the changes in T cell polarity over time. PBMCs were stimulated with a SUDV GP-specific peptide pool and analyzed for CD4^+^ effector memory re-expressing (EM-RE) T cells (Fig. S4). Levels of CD69, IFNγ and IL-4 expression were analyzed at the time of vaccination, challenge, and at selected exam days after challenge for all groups. In addition, d-14 samples were analyzed for the control, VSV-EBOV and VSV-SUDV d-28 groups. No significant differences were observed between any groups at all time points. However, upon further investigation, a shift was observed in the T cell polarity of the antigen-specific CD4^+^ EM-RE cells for the VSV-SUDV groups (Fig. S4). Initially, this memory cell cohort was primarily Th1-driven in nature expressing IFNγ at challenge (d0). As the cellular immunity memory matured over time, the CD4^+^ EM-RE cells shifted to a balanced Th1/Th2 phenotype expressing either IFNγ or IL-4 likely contributing to the maintenance of the humoral immune response.

## Discussion

Uganda declared its 6^th^ outbreak of SVD on January 30^th^ 2025 and to date has reported 14 cases (12 confirmed, 2 probable) with 4 fatalities (2 confirmed, 2 probable) ^5^. Despite the lack of approved vaccines, several vaccine candidates ^26^ have demonstrated efficacy in preclinical NHP studies including a VSV-SUDV vaccine following the success of the related EBOV vaccine, VSV-EBOV (Ervebo by Merck). This vaccine is in clinical development by IAVI, has shown single dose efficacy in NHPs within 4 weeks and immunogenicity in a Phase I Clinical trial ^27^. Uganda started a ring vaccination trial with this vaccine in February 2025 ^6^. However, the fast-acting potential of this vaccine has not been investigated.

We previously updated VSV-SUDV by using a more current SUDV GP from the SUDV-Gulu strain and demonstrated uniform protection with a single dose within 4 weeks in NHPs with preexisting EBOV immunity ^9^. The current study addresses two main knowledge gaps, the rapid onset of protection with a single dose of the VSV-SUDV vaccine and the need for species-specific vaccines during outbreaks as the VSV-EBOV vaccine did not provide any protective benefit against lethal SUDV challenge. Single-dose VSV-SUDV vaccination 28 or 7 days before lethal SUDV challenge protected naïve NHPs from disease and lethality independent of vaccination time point highlighting the fast-acting potential of the VSV-SUDV vaccine. In addition, we demonstrate the lack of cross-protection from VSV-EBOV vaccine against SUDV infection despite the many similarities between the viruses including the development of cross-reactive IgG responses after vaccination. This contrasts with a recent study exploring cross-protection of VSV-EBOV against SUDV in the guinea pig disease model which resulted in ∼60% survival ^28^. However, the NHP model is regarded as the “gold standard” in filovirus countermeasure research and efficacy results seem more predictive for human outcome than results from rodent challenge studies ^29,30^. This highlights the continued need for NHP studies on filovirus countermeasure development and approval following the animal rule ^31^.

In a previous study we showed that VSV-EBOV vaccination ∼1 year before SUDV challenge only resulted in limited cross-reactivity but no protective efficacy ^9^. Here, we provide additional proof for our previous finding of no benefit from VSV-EBOV vaccination against SUDV challenge. Furthermore, our results indicate that the time between VSV-EBOV vaccination and SUDV exposure may impact the disease course as the VSV-EBOV cohort succumbed significantly earlier compared to the control group vaccinated with VSV-LASV, yet disease severity was the same. VSV-LASV has been shown in the past to not provide any cross-protection against filovirus infection in NHPs ^12^. We examined serum antibodies for infection-enhancing properties but could only detect background levels of ∼4% of GFP-positive cells which are negligible in comparison to previous studies describing filovirus ADE using monoclonal antibodies at ∼60% ^24^. Similarly, for Dengue ADE levels are much higher at 20-40% ^32^ leading us to believe that ADE is not the primary mechanism for the differences in disease progression we observed after SUDV challenge. A secondary mechanism for ADE of disease is through the binding of the complement C1q receptor ^22,33^. However, we found no significant increase in complement deposition for the VSV-EBOV vaccinated NHPs compared to the controls (Fig. 5F). One may speculate that 4 weeks after vaccination is likely the time when the immune response specific to EBOV GP is at high and very strong, yet unspecific responses elicited by VSV-EBOV may accelerate disease. We have yet to determine why this accelerated disease was observed 28 days after vaccination but not one year after VSV-EBOV vaccination and EBOV challenge ^9^.

The VSV-SUDV group vaccinated on d-7 demonstrated no outward signs of disease similar to the group vaccinated with VSV-SUDV on d-28, however, only the d-7 group did not seem to have an impact on levels of platelets, calcium or certain cytokines including IL-6 and TNFα. Notably, the VSV-EBOV-vaccinated NHPs developed lung pathology after SUDV infection which was less severe in control NHPs at the time of necropsy. Cytokine analysis of lung homogenates and serum revealed high levels of the pro-inflammatory cytokine IL-6 in this group likely contributing to liver damage and the loss of albumin associated with high levels of IL-6 ^34^. Other highly expressed cytokines and chemokines are GM-CSF, TNFα, IP-10, and MCP-1 which are associated with the dysregulated inflammatory response, particularly macrophage activation. Another significant difference between the VSV-EBOV and control groups is the neutrophilia observed on d3 for the VSV-EBOV group only. The induction of aberrant neutrophils early after SUDV challenge has previously been described as a contributing factor to fatal disease and may have had the same effect here ^35,36^. Collectively, our data indicates that an aberrant innate immune response in expanded tissue compartments early in the disease course may contribute to the accelerated time to death of the VSV-EBOV-vaccinated NHPs.

While we measured significant levels of EBOV GP-specific IgG cross-reactive to SUDV GP in VSV-EBOV-vaccinated NHPs, their neutralization capacity was limited. This finding is in line with a recent publication describing limited to no neutralizing activity against SUDV in serum of humans vaccinated with a single dose of VSV-EBOV ^37^. This lack of cross-neutralization was overcome by a boost vaccination with VSV-EBOV; however, it is unclear if the level of neutralization would provide any benefit after a SUDV infection ^37^. In our previous study NHPs vaccinated with VSV-EBOV and challenged with EBOV were not protected from SUDV challenge one year later suggesting that cross-reactive immunity may wane over time ^9^. As our study assessed protection after a single dose vaccination only, future studies targeted towards cross-protective efficacy will investigate this further.

Cross-protection among ebolaviruses and MARV can be achieved with a single VSV-based filovirus vaccine ^38^, a mixture of individual VSV-based vectors ^12^ or prime/boost of a cocktail of filovirus GPs ^39^ indicating that the presence of the antigen specific to the challenge virus may be important. However, the mixture of individual VSV-based vectors provided protection against Taï forest virus which was not part of the antigen cocktail ^12^ challenging this hypothesis. In addition, a more recent study by van Tol and colleagues showed that single dose as well as prime/boost vaccination with an adenovirus-based vector encoding both the EBOV GP and the SUDV GP did not result in protection from SUDV challenge ^36^. Like the data presented here for the VSV-EBOV group, all NHPs vaccinated with this adenovirus-based vector had cross-reactive binding antibodies but failed to protect from SUDV challenge. The neutralizing response was much more potent against EBOV than SUDV. At this point it is unclear if the vaccine would have protected against EBOV. Taken together, it seems that there is an importance to the specificity of the protective immune response at least against infection with ebolaviruses that warrants further investigation.

Our further inquiry into Fc effector functions revealed that ADCP is stimulated when vaccination is administered longer than 7 days prior to challenge; however, there was no significant difference between the d-7 cohort that survived the infection and the VSV-EBOV cohort which succumbed to disease indicating that ADCP is not a correlate of protection. While myeloid cells can be the target of early viral replication these phagocytotic cells have the capacity to degrade the virions which they engulf. This is apparent in the ADNP data demonstrating that both vaccine cohorts have significantly higher induction of neutrophil phagocytosis compared to the control and VSV-EBOV cohorts on d6. The addition of the antibody immune complex upon internalization may lock the virions in conformation inhibiting viral receptor binding upon maturation of the endosome ^40^. Typically, binding to the Fc receptor initiates downstream signaling to promote antiviral functions ^40^. In our study this was not the case for the potential protective role of the early innate lymphocytes, in which we saw little activation of ADNKA.

In addition, both VSV-SUDV vaccine cohorts demonstrated significantly elevated levels of C3 deposition indicative of ADCD compared to the control and VSV-EBOV cohorts at d6, the time when the control and VSV-EBOV groups succumbed to disease. The role in complement activation in filovirus pathogenesis until this point is a double-edged sword. Overactivation of the complement pathway via the classical route of activation and mannose binding lectin route have contributed to increased pathogenesis ^22,41,42^. However, in several monoclonal antibody treatment studies it has been demonstrated that the engineering of antibodies to induce ADCD increased their therapeutic potential ^43,44^. Aligning with our data NHPs in the VSV-SUDV cohorts had significantly higher C3 deposition on d6 but did not succumb to disease like the control and VSV-EBOV cohorts which did not demonstrate high levels of ADCD at that time. In summation the antibody functionality profile indicates that vaccinated animals which develop ADNP and ADCD survived the challenge, and animals vaccinated at a longer interval from challenge induced a stronger ADCP. These results indicate that a diverse antibody functionality repertoire is required to inhibit disease progression after SUDV infection.

It has previously been established that the primary immune mechanism of protection of VSV vaccination against filoviruses is the humoral response ^19,20^. Therefore, we sought to investigate the cellular response for supportive roles to aid in the development of the humoral response ^25^. We did not determine differences between the vaccinated NHPs and the control and VSV-EBOV NHPs at a time of peak disease (d6). Upon longitudinal investigation of the CD4 T cell response, we observed a shift in the T cell polarity of the antigen specific CD4^+^ EM-RE cells. Initially this memory cell cohort is primarily Th1 in nature expressing IFNγ. As the cellular immunity memory matures the CD4^+^ EM-RE T cells shift to a balanced Th1/Th2 expressing either IFNγ or IL-4.

This is important in the maintenance of the humoral response as IL-4 activates mature B cells for antibody secretion, as well as promotes the survival of B cells ^45,46^. This balance during maturation contributes to a robust vaccination response, in both direct antiviral effector functions of CD4^+^ EM-RE Th1 T cells as well as the support of the humoral response via the CD4^+^ EM-RE Th2 T cells. In addition to the total antigen-specific humoral response and the neutralization capacity of that response, alternative Fc functions are induced.

While this study provides insight into the protective immunity elicited by the VSV-SUDV vaccine, it also presents its limitations regarding the investigation of cross-protection. If d-7 group vaccinated with VSV-EBOV would have been included, a more thorough investigation of the timing for the onset of a potentially cross-protective immune response could have been performed. Likewise, a control group vaccinated on d-7 was also not included, however, the impact of the data from a VSV-EBOV d-7 group would have likely been greater. Furthermore, T cell responses and ADE could have been investigated in more detail, however, the sampling time points and amounts of samples collected limited us in pursuing this. Future studies investigating cross-protection between filoviruses will be designed with additional analyses in mind. Lastly, we chose the IM challenge method as it had been used in our previous studies, however, future studies may be conducted using mucosal challenge routes to mimic natural infection closer. At this time, however, a mucosal challenge for NHPs has not been established at our facility.

Taken together, the data described herein demonstrates that a single vaccination with VSV-SUDV is 100% protective against SUDV challenge in the cynomolgus macaque model. Moreover, we also demonstrate that the VSV-SUDV vaccine provides complete protection when administered only 7 days prior to challenge. A fast-acting SUDV vaccine is highly advantageous for quelling SUDV outbreaks such as the current one in Uganda ^5^. In fact, a VSV-SUDV vaccine expressing a slightly different SUDV GP supported through the International AIDS Vaccine Initiative (IAVI) is currently being deployed in Uganda in a clinical ring-vaccination trial ^6^. This is reminiscent of the situation in West Africa during the Ebola virus disease outbreak where concomitantly VSV-EBOV was shown to possess fast-acting potential as an emergency vaccine ^15^. Thus, our study supports the started ring vaccination trial from a preclinical point of view. In addition, our study here clearly shows that species-specific VSV vaccines are needed for control measures and provides preclinical evidence for potential negative effects of using an unmatched VSV vectored vaccine. While there is a push towards pan-filovirus protective countermeasures particularly for frontline workers, more work is needed to support that concept.

## Materials & Methods

### Ethics statement

All work involving SUDV was performed in the maximum containment laboratory (MCL) at the Rocky Mountain Laboratories (RML), Division of Intramural Research, National Institute of Allergy and Infectious Diseases, National Institutes of Health. RML is an AAALAC International-accredited institution. All procedures followed RML Institutional Biosafety Committee (IBC)-approved standard operating procedures (SOPs). Animal work was performed in strict accordance with the recommendations described in the Guide for the Care and Use of Laboratory Animals of the National Institute of Health, the Office of Animal Welfare and the Animal Welfare Act, United States Department of Agriculture. This study was approved by the RML Animal Care and Use Committee (ACUC), and all procedures were conducted on anesthetized animals by trained personnel under the supervision of board-certified clinical veterinarians. The animals were observed at least twice daily for clinical signs of disease according to a RML ACUC-approved scoring sheet and humanely euthanized when they reached endpoint criteria.

Animals were housed in adjoining individual primate cages that enabled social interactions, under controlled conditions of humidity, temperature, and light (12 hours light - dark cycles). Food and water were available *ad libitum*. Animals were monitored and fed commercial monkey chow, treats, and fruit at least twice a day by trained personnel. Environmental enrichment consisted of commercial toys, music, video and social interaction.

### Cells, vaccines, and challenge virus

Vero E6 cells (*Mycoplasma* negative; ATCC, Cat. No. CRL-1586) were grown at 37°C and 5% CO_2_ in Dulbecco’s modified Eagle’s medium (DMEM) (Sigma-Aldrich, St. Louis, MO) containing 10% fetal bovine serum (FBS) (Wisent Inc., St. Bruno, Canada), 2 mM L-glutamine, 50 U/mL penicillin, and 50 mg/mL streptomycin (all supplements from Thermo Fisher Scientific, Waltham, MA). THP-1 (*Mycoplasma* negative) were grown at 37°C and 5% CO_2_ in Roswell Park Memorial Institute (RPMI) medium (Sigma-Aldrich) containing 10% FBS, 2 mM L-glutamine, 50 U/mL penicillin, and 50 mg/mL. HL-60 cells (*Mycoplasma* negative; ATCC; Cat. No. CCL-240) were propagated in Iscove’s Modified Dulbecco’s medium (IMDM; Sigma-Aldrich) with 20% FBS, 2 mM L-glutamine, 50 U/mL penicillin, and 50 mg/mL streptomycin and differentiated with 1.3% DMSO for 5 days. NK92 cells expressing CD16 (*Mycoplasma* negative; ATCC Cat No. PTA-6967) were propagated in minimal essential medium (MEM; Sigma-Aldrich) supplemented with 12.5% FBS, 12.5% horse serum, 10µg human IL-2, 2 mM L-glutamine, 50 U/mL penicillin, and 50 mg/mL streptomycin. K562 cells (*Mycoplasma* negative; ATCC Cat. No. CCL-243) were propagated in IMDM containing 10% FBS, 2 mM L-glutamine, 50 U/mL penicillin, and 50 mg/mL streptomycin. Previously described VSV-based vaccine vectors expressing the EBOV-Kikwit GP (VSV-EBOV)^15^, SUDV-Gulu GP (VSV-SUDV)^9^, or Lassa virus GPC (VSV-LASV) ^47^ were used in this study. For the NHP challenge, SUDV-Gulu (GenBank NC_006432.1) at 10,000 TDCID_50_ was used IM as previously described ^9^.

### NHP study design

Twenty-four male cynomolgus macaques of Chinese or Cambodian origin, 2.5-4.5 years of age and 2.9-5.2 kg in weight at the time of vaccination, were used in this study. The NHPs were randomly divided into 4 study groups with 6 NHPs each. On d-28, NHPs received a 1 ml IM injection for vaccination into 2 sites in the caudal thighs containing either 1×10^7^ PFU VSV-SUDV (n=6), 1x 10^7^ PFU VSV-EBOV (n=6) or 1x 10^7^ PFU VSV-LASV (control; n=6). After vaccination, physical examinations and blood draws were performed on d-27, −25, −21, and −14. On d-7 the last vaccine group received 1×10^7^ PFU VSV-SUDV IM (n=6). For this group, physical examinations and blood draws were performed on d-6, and −4. All 24 NHPs were challenged IM on d0 with 1×10^4^ TCID_50_ SUDV-Gulu (backtitered as 1.1×10^4^ TCID_50_ per NHP) into 2 sites in the caudal thighs as previously described ^9^. Physical examinations and blood draws were performed on d0, 3, 6, 9, 14, 21, 28, and 35 and at euthanasia (d42 for survivors; humane endpoint for non-survivors). Following euthanasia, a necropsy was performed, and tissue samples were collected for analysis.

### Hematology and serum chemistries

Complete blood cell counts were determined from EDTA blood with the IDEXX ProCyte DX analyzer (IDEXX Laboratories, Westbrook, ME). Serum biochemistry was analyzed with a Vetscan 2 using Preventive care profile disks (Abaxis, Union City, CA).

### VSV and SUDV RNA

VSV RNA copy numbers in EDTA blood samples after vaccination were determined as previously described ^48^. SUDV RNA copy numbers in EDTA blood and tissue samples after challenge were determined as previously described ^9^.

### Histology and immunohistochemistry

Necropsies and tissue sampling were performed according to IBC-approved SOPs. Collected tissues were fixed for 8 days in 10% neutral-buffered formalin, embedded in paraffin, processed using a VIP-6 Tissue Tek (Sakura Finetek USA, Torrance, CA) tissue processor, and embedded in Ultraffin paraffin polymer (Cancer Diagnostics, Durham, NC). Samples were sectioned at 5 μm, dried overnight at 42°C, and resulting slides were stained with hematoxylin and eosin. Specific anti-VP40 immunoreactivity was detected using a previously described cross-reactive rabbit anti-EBOV VP40 polyclonal antibody (a generous gift by Dr. Yoshihiro Kawaoka, University of Wisconsin-Madison) at a 1:1000 dilution. The immunohistochemical assay was carried out on a Discovery ULTRA automated staining instrument (Roche Tissue Diagnostics, UK) with a Discovery ChromoMap DAB (Ventana Medical Systems, Oro Valley, AZ) kit. All tissue slides were evaluated by a board-certified veterinary pathologist.

### Assessment of humoral immune response

Post-challenge NHP sera were inactivated by γ-irradiation (4 MRad), a well-established method with minimal impact on serum antibody binding ^49^ and removed from the MCL according to SOPs approved by the RML IBC. Titers for IgG specific to EBOV GP or SUDV GP were determined in serum samples using ELISA kits following manufacturer’s instructions (Alpha Diagnostics, San Antonio, TX). Serum samples collected between d-28 and d0 were diluted 1:200; samples collected d3 and later in the study were assessed at 1:1,000 dilution.

Neutralization assay with VSV-SUDV-GFP was performed as previously described ^9^. The VSV-SUDV-GFP assay was optimized so incubation of the serum/virus mix on the Vero E6 cells lasted for 16 hours at 37°C. Samples were run on the FACSymphony A5 Cell Analyzer (BD Biosciences, Mississauga, ON, Canada) and the GFP-positive cell count was determined. Gating strategy is shown in Fig. S6A.

### SUDV sGP ELISA

The SUDV sGP-specific sandwich ELISA was performed as previously described with minor differences ^18^. SUDV sGP in serum and homogenized, cleared, γ-irradiated (4 MRad) lung samples at 1:100 dilution was captured using 1ug/ml polyclonal rabbit anti-SUDV sGP antibody (IBT Bioservices, Rockville, MD; Cat. No. 0302-030). Dilutions of recombinant SUDV sGP (IBT Bioservices, Cat. No. 0570-001) at known concentrations served as standards.

### Antibody-dependent enhancement (ADE) assay

K562 cells (*Mycoplasma* negative; ATCC Cat. No. CCL-243) were used to determine if serum antibodies could enhance SUDV infection from a protocol adapted from ^32^. Briefly, NHP serum was diluted 1:10 and thereafter threefold and then incubated with SUDV-GFP ^37^ at a multiplicity of infection (MOI) of 0.5 for 1 h at 37 °C. Approximately 1 × 10^5^ K562 cells were added to each well in a 96 well-plate containing the mixture of the virus and the serum and incubated at 37 °C for 48 hours. Then, cells were washed and fixed with 4% paraformaldehyde (PFA). Data were acquired on a Cytoflex LX (Beckman Coulter, Brea, CA) and analyzed in FlowJo v10. Gating strategy is shown in Fig. S5. Percent enhanced infection was determined by subtracting the GFP signal of the viral control (no serum) to the treated samples.

### Quantification of Antibody Effector Functions

Assays for antibody effector functions were adapted from previously established protocols ^50,51^. Recombinant SUDV GPΔTM (IBT Bioservices) was tethered to Fluospheres NutrAvidin-Microspheres yellow-green or red (Thermo Fisher Scientific) using the EZ-link Micro Sulfo-NHS-LC-Biotinylation kit (Thermo Fisher Scientific).

#### ADCD

Serum samples were heat-inactivated at 56°C for 30 min then diluted 1:200 in DMEM and applied to the conjugated beads (20 μl/well) for one hour at 37°C. Next, guinea pig complement (Cedarlane, Burlington, Canada) was added for 30 minutes. The bead complexes were washed with PBS containing 15 mM EDTA and stained with anti-C3c-FITC (Antibodies-Online, Pottstown, PA; Cat. No. ABIN458597). Data were acquired on a FACS Symphony (BD Biosciences, Franklin Lakes, NJ) and analyzed in FlowJo v10. Gating strategy is shown in Fig. S6B.

#### ADNP

Biotinylated SUDV GPΔTM was coupled to yellow-green Neutravidin beads (Life Technologies). Serum samples were diluted 1:250 in culture medium and incubated with GP-coated beads for 2 hours at 37 °C. Beads (20 μl) were added to HL-60 cells (5×10^4^ cells/well) and incubated for 2 hours at 37 °C. Cells were then stained for CD11b (Clone G10F5; BioLegend), CD16 (Clone UCHT1; BD Biosciences), and fixed with 4% paraformaldehyde. Data were acquired on a FACS Symphony (BD) and analyzed in FlowJo v10. Neutrophils were defined as SSC-A^high^ CD11b^+^, CD16+. A phagocytic score was determined using the following formula: (percentage of FITC^+^ cells)*(geometric mean fluorescent intensity (gMFI) of the FITC^+^ cells)/10,000. Gating strategy is shown in Fig. S6D.

#### ADCP

Serum samples were diluted 1:250 in DMEM and incubated with 20 μl of the conjugated beads for 2 hours at 37°C. The serum/bead mixture was then transferred to a plate of THP-1 cells (2.5×10^4^ cells/well) and incubated overnight at 37°C. Cells were fixed with 4% paraformaldehyde. Data were acquired on a FACS Symphony (BD) and analyzed in FlowJo v10. Gating strategy is shown in Fig. S6C. A phagocytic score was determined using the following formula: (percentage of FITC^+^ cells)*(geometric mean fluorescent intensity (gMFI) of the FITC^+^ cells)/10,000.

#### ADNKA

Recombinant SUDV GPΔTM (IBT Bioservices) was coated onto MaxiSorp 96-well plates (Thermo Fisher Scientific) at 300 ng/well at 4 °C overnight. Wells were washed with PBS and blocked with 5% BSA prior to addition of serum (1:100 dilution) and incubation for 2 hours at 37 °C. Unbound antibodies were removed by washing with PBS, and NK92 cells expressing CD16 were added at 5×10^4^ cells/well in the presence of 4 μg/ml brefeldin A (Sigma-Aldrich), 5 μg/ml GolgiStop (Thermo Fisher Scientific) and anti-CD107a antibody (Clone H4A3, BD Biosciences) for 5 hours. Cells were surface stained for CD16 (clone 3G8, Pacific Blue; Biolegend Cat. No. 302032) and CD56 (clone 5.1H11; Alexa Fluor488; Biolegend Cat. No. 362518) for 20 min. Cells were fixed and permeabilized with Fix/Perm (Life Technologies) according to the manufacturer’s instructions to stain for intracellular IFNγ (Clone B27, PE; Biolegend Cat. No. 506507) and TNFα (clone Mab11, APC; Biolegend Cat. No. 502912). Data were acquired on a FACS Symphony (BD Biosciences) and analyzed in FlowJo v10. Gating strategy is shown in Fig. S6E.

### Cellular Phenotyping Assays

PBMCs were isolated from whole blood samples, stored, and revived as previously described ^51^. For T cell response analysis, PBMCs were stimulated with 1.5 μg/ml of either a SUDV GP peptide pool, media alone, or a SARS-CoV-2 nucleocapsid peptide pool as an unspecific control for 16 hours. Initial peptide pool activation media was then removed and 1.5 μg/ml of the SUDV GP peptide pool, media alone, or the SARS-CoV-2 nucleocapsid peptide pool together with 2.5μg/ml Brefeldin A (Biolegend) cell stimulation cocktail (containing PMA-Ionomycin, Biolegend) was added to the cells for 5 hours. Following, cells were surface-stained and analyzed as previously described ^51^. NK cell immune responses were measured as previously described ^51^. Gating strategy is shown in Fig. S7.

### Cytokine and chemokine analysis

Cleared lung homogenates and serum samples were inactivated by γ-irradiation. Levels of GM-CSF, IFN-γ, IFN-α2a, IL-1β, IL-4, IL-6, IL-10, IL-12p70, IL-15, IP-10, MCP-1, and TNF-α were assessed using a customized Meso Scale Discovery (MSD; Rockville, MD, USA) U-PLEX NHP multiplex assay according to manufacturer’s instructions. Samples were read on the MSD MESO QuickPlex SQ 120MM with Methodical Mind software (v. 1.0.38) and analyzed with MSD Discovery Workbench software (v. 4.0).

### Statistical analysis

Statistical analysis was performed in Prism 9 (GraphPad). For statistical analysis, data from the VSV-EBOV group on d5 were combined with the single d6 data point and compared to all other groups on d6. Most data was analyzed by two-way ANOVA with Tukey’s multiple comparison to evaluate statistical significance at all timepoints between all groups. Data depicted in Fig. S5A was analyzed by Kruskal-Wallis test with Dunn’s multiple comparisons. Data in Fig. S5B was analyzed by Mann-Whitney test. Significant differences in the survival curves shown in Fig. 1C were determined performing Log-Rank analysis. Statistical significance is indicated as p<0.0001 (****), p<0.001 (***), p<0.01 (**), and *p*<0.05 (*).

## Author Contributions

HF and AM conceived the idea, and designed the study. BMG, HF and AM secured funding. PF, FF, BJS, and AM conducted the NHP study. PF, KLO, JFR, CAP, CSC, BMG, and AM processed samples, performed assays, and analyzed the data. AM wrote the manuscript with input from all authors. All authors had full access to all the data in the study and had final responsibility for the decision to submit for publication.

## Acknowledgments

We thank members of the Rocky Mountain Veterinary Branch, NIAID for supporting this study. We are grateful for Dr. C. Albarino who shared the SUDV-GFP with us under a material transfer agreement. We also thank Dr. James D. Brien, University of Kentucky, for his assistance establishing the ADE assay.

## Conflict of Interests

HF claims intellectual property of VSV-based filovirus vaccines. All other authors declare no conflicts of interest.

## Funding

The study was funded by the Intramural Research Program, NIAID, NIH. Funding was also provided in part by NIH/NIAID U01-AI151799 and R01-AI175698 to BMG. The funders had no role in study design, study conduct, data analysis or decision to publish.

## Data availability

All data is presented in the paper and supplemental material and available at 10.6084/m9.figshare.28661588.

---

**Figure S1.**
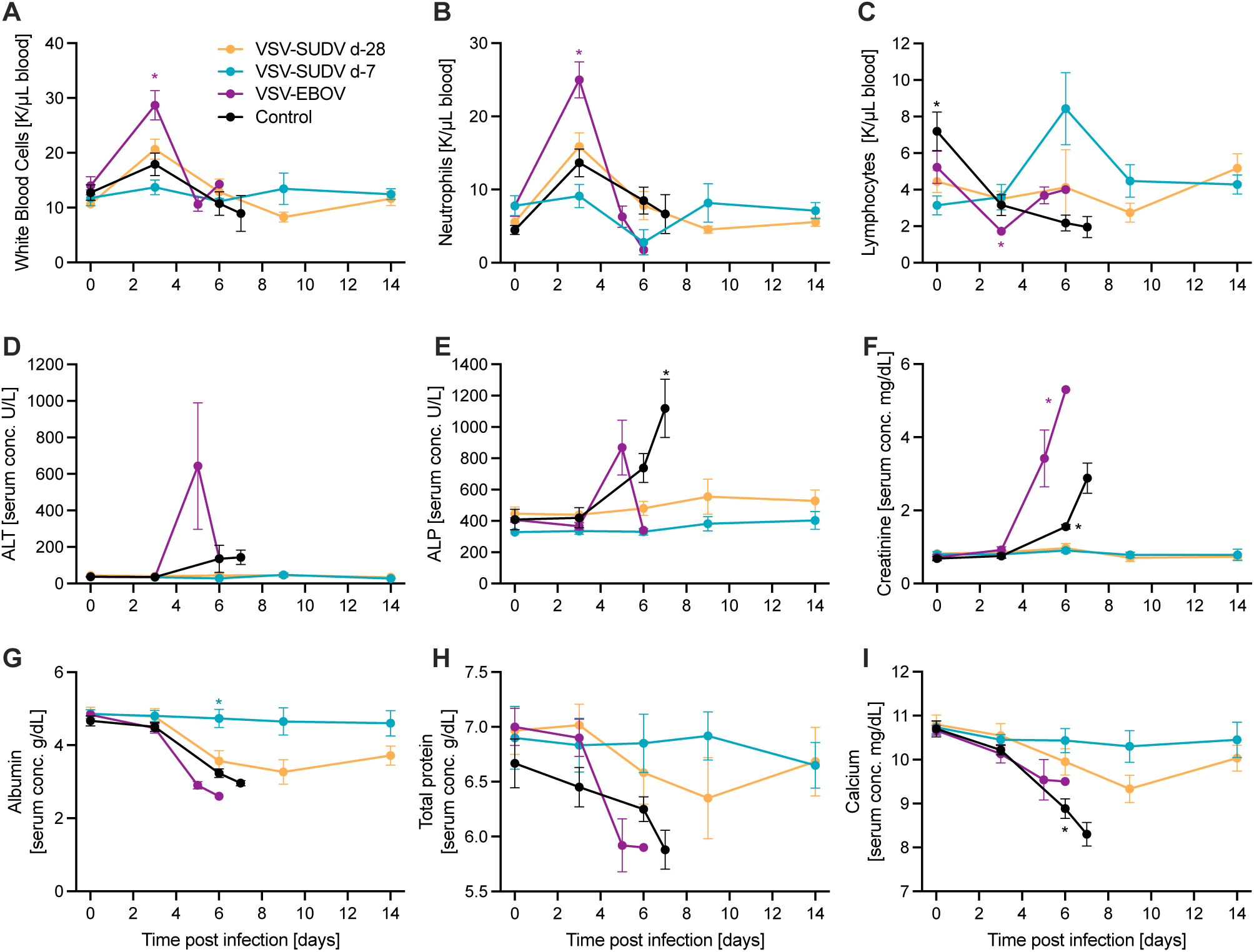
Changes in blood and serum parameters in NHPs after SUDV challenge. NHPs (n=6 per group) were vaccinated 28 days before challenge (d-28) with either VSV-SUDV, VSV-EBOV or control vaccine (VSV-LASV). Another group was vaccinated with VSV-SUDV 7 days before challenge (d-7). On day 0, all 24 NHPs were challenged with a lethal dose of SUDV. Changes in (A) white blood cell, (B) neutrophil, and (C) lymphocyte counts as well as serum levels of (D) alanine aminotransferase (ALT), (E) alanine phosphatase (ALP), (F) creatinine, (G) albumin, (H) total protein and (I) calcium are shown. Statistical significance was determined by two-way ANOVA with Tukey’s multiple comparisons. Statistical significance is indicated as \**p*<0.05.

**Figure S2.**
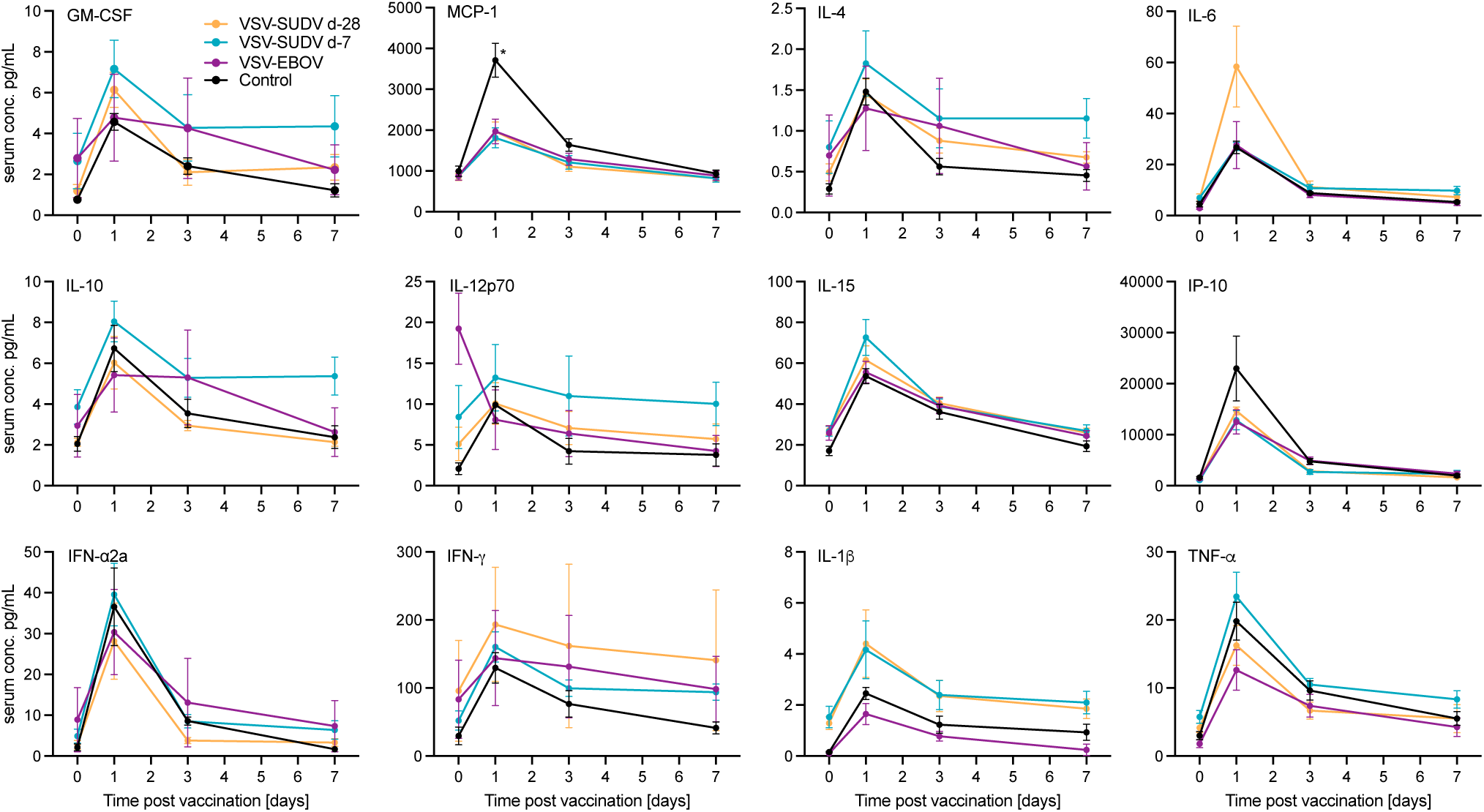
Levels of cytokines and chemokines in the serum after vaccination. Expression levels of selected cytokines or chemokines were determined after vaccination. Mean and standard error of the mean are depicted. Statistical significance was determined by two-way ANOVA with Tukey’s multiple comparisons. Statistical significance is indicated as \**p*<0.05.

**Figure S3.**
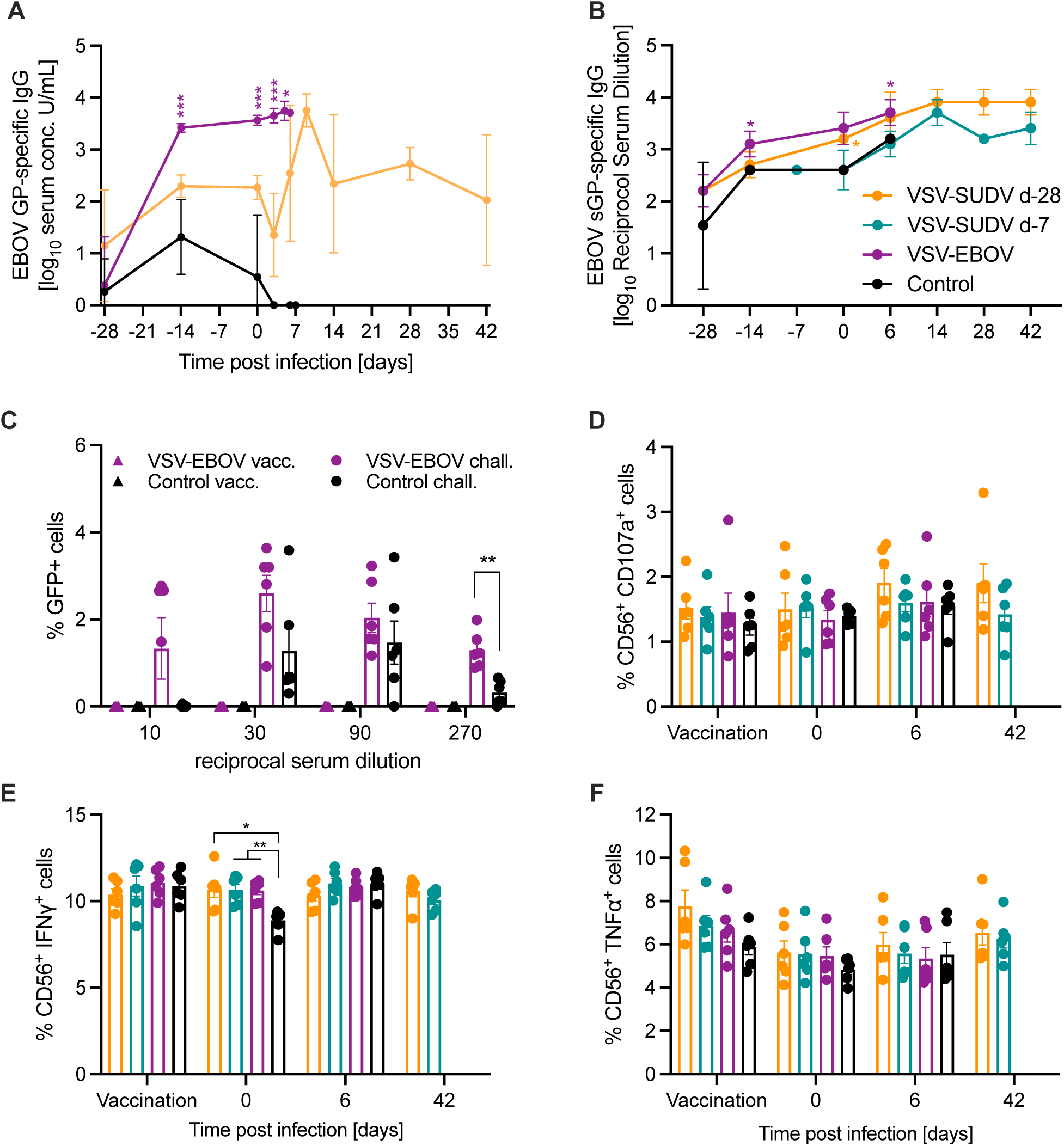
EBOV GP/sGP-specific IgG levels, antibody-dependent enhancement (ADE) and antibody-dependent killer cell activation (ADNKA). (A) EBOV GP-specific IgG responses in the serum of the 3 groups vaccinated 28 days before SUDV challenge. (B) EBOV sGP-specific IgG responses over time. (C) ADE on K562 cells with SUDV-GFP. ADNKA with (D) CD107^+^ cells, (E) IFNγ^+^ cells, and (F) TNFα^+^ cells. Geometric mean and geometric SD are depicted in A, B. Mean and standard error of the mean are shown in C-F. Statistical significance was determined by two-way ANOVA with Tukey’s multiple comparisons (3 or 4 groups) or Sidak’s multiple comparisons (2 groups). Statistical significance is indicated as \**p*<0.05, \*\**p*<0.01, and \*\*\**p*<0.001.

**Figure S4.**
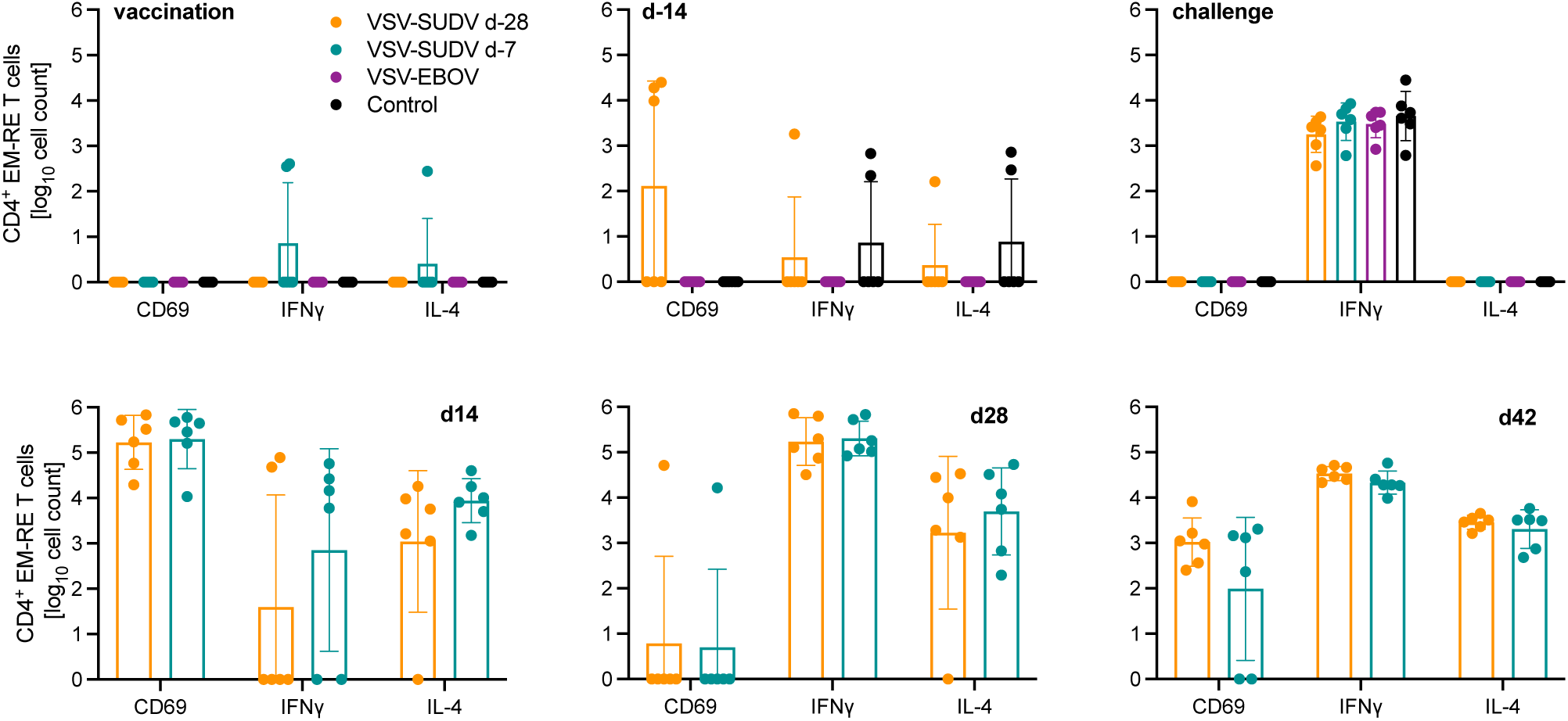
CD4 T cell responses after vaccination and challenge. PBMCs were stimulated with a SUDV GP-specific peptide pool and analyzed for CD4^+^ effector memory re-expressing (EM-RE) T cells. Levels of CD69, IFNγ and IL-4 expression were analyzed throughout the study. Geometric mean and geometric SD are depicted. Data was analyzed by two-way ANOVA with Tukey’s multiple comparisons and no statistical significance was determined between groups. Neutralization was assessed in Vero E6 cells with VSV-SUDV-GFP. ADE was assessed using K-562 cells and SUDV-GFP. GFP-positive cells were counted. Exemplary gating strategy for ADE with serum from a (A) VSV-EBOV-vaccinated NHP and (B) control NHP at the time of SUDV challenge.

**Figure S5.**
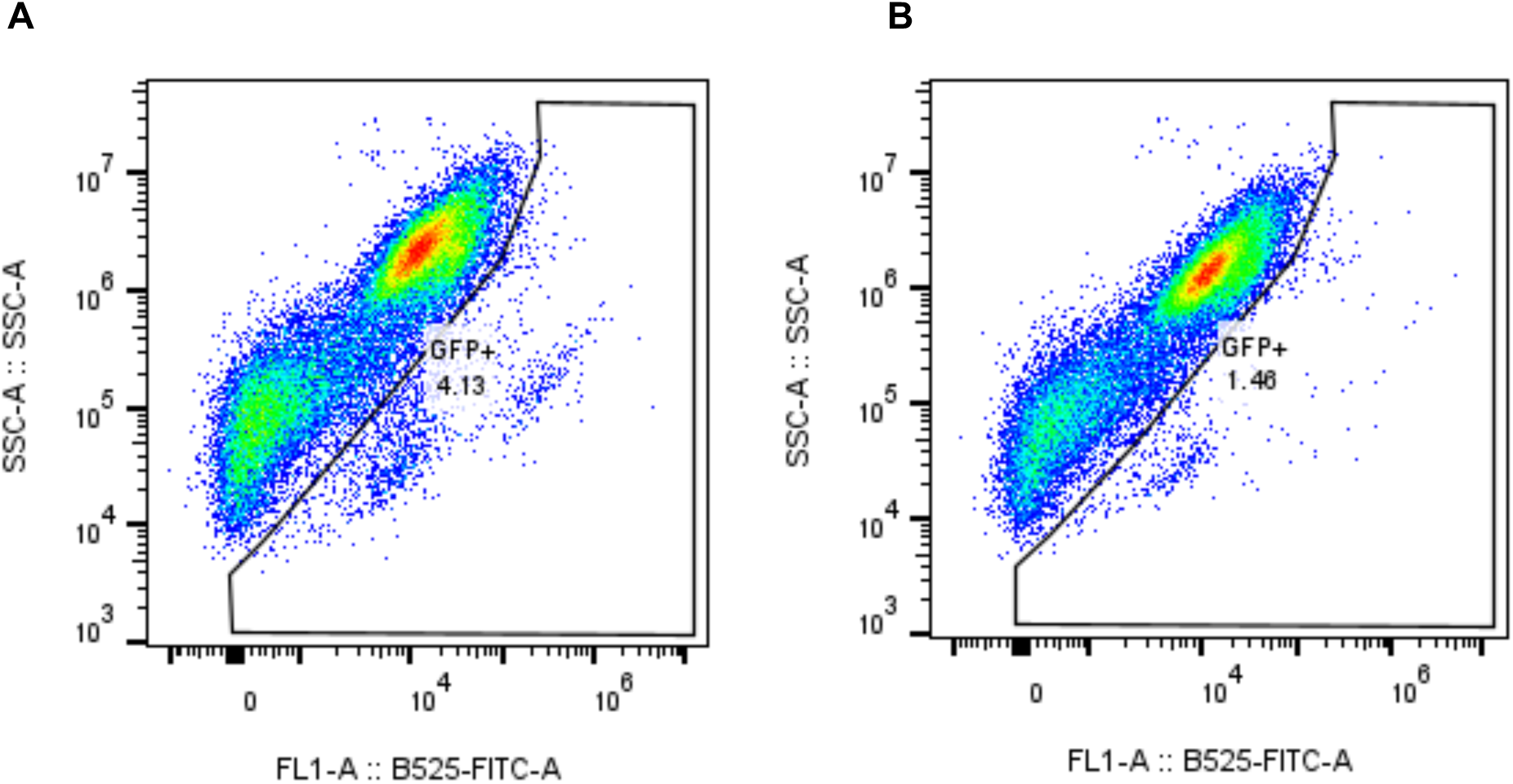
Gating strategy for neutralization and antibody-dependent enhancement (ADE) assays. Neutralization was assessed in Vero E6 cells with VSV-SUDV-GFP. ADE was assessed using K-562 cells and SUDV-GFP. GFP-positive cells were counted. Exemplary gating strategy for ADE with serum from a (A) VSV-EBOV-vaccinated NHP and (B) control NHP at the time of SUDV challenge.

**Figure S6.**
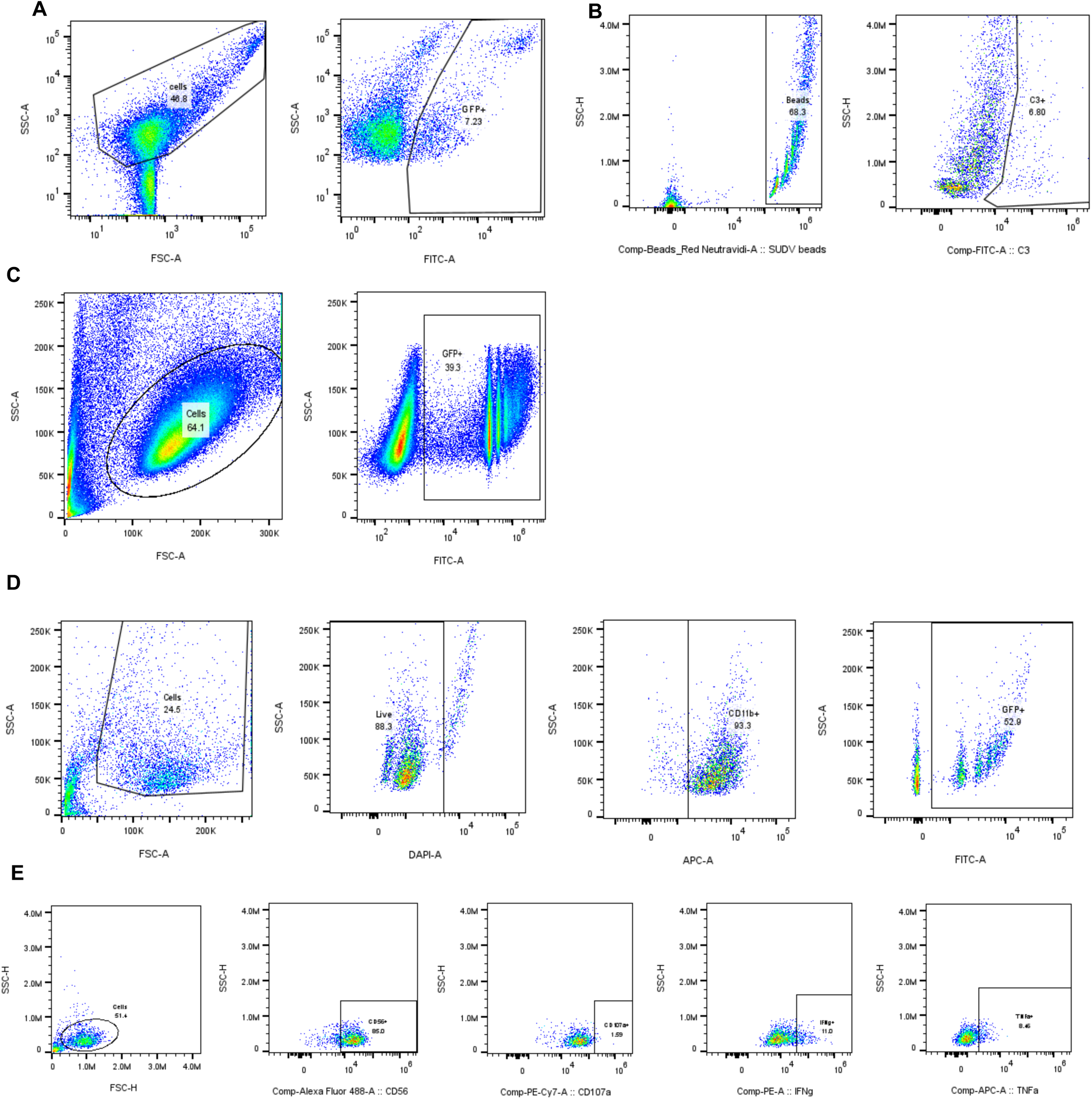
Gating strategy for antibody effector function analysis. (A) Neutralization. (B) Antibody-dependent complement deposition (ADCD). (C) Antibody-dependent cellular phagocytosis (ADCP). (D) Antibody-dependent neutrophil phagocytosis (ADNP). (E) Antibody-dependent natural killer cell activation (ADNKA).

**Figure S7.**
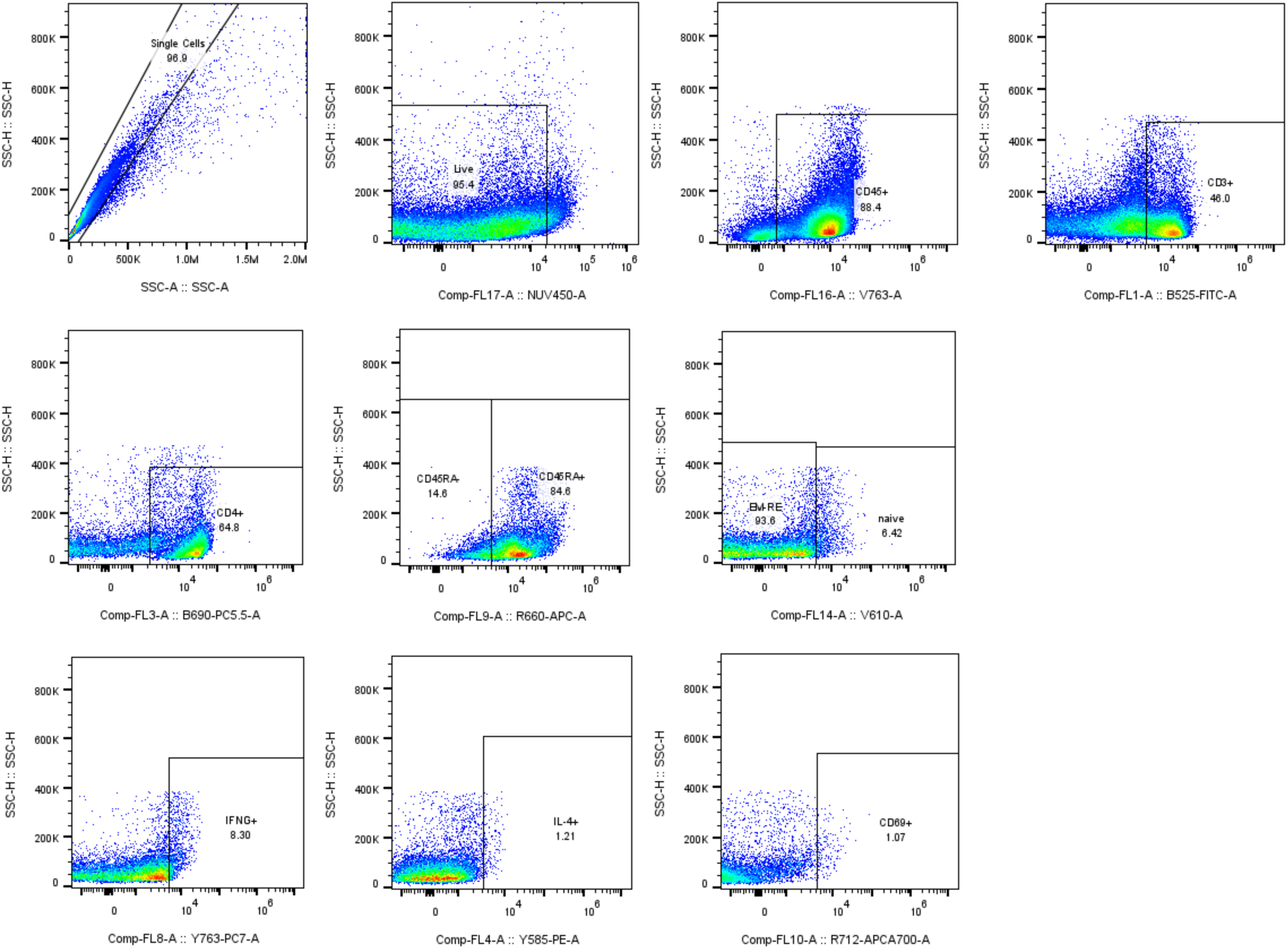
Gating strategy for functional analysis of vaccine-elicited T cells. Frozen PBMCs were thawed and restimulated overnight with a SUDV GP-specific peptide pool.

